# Identification of MED13 and DDX60 as critical host factors for SARS-CoV-2 infections

**DOI:** 10.64898/2026.02.25.707691

**Authors:** Hyesoo Kwon, Sha Tim Wai, Georg Michlits, Matheus Dyczynski, Ana Marković, Andreia Sofia Batista Rocha Berger, Lijo John, Moritz Horn, Friedemann Weber, Josef M. Penninger, Vanessa M. Monteil, Ali Mirazimi

**Affiliations:** Swedish Veterinary Agency, Uppsala, Sweden; Karolinska Institute and Karolinska University Hospital, Department of Laboratory Medicine, Unit of Clinical Microbiology, Stockholm, Sweden; Institute for Virology, FB10-Veterinary Medicine, Justus-Liebig University, Giessen, Germany; JLP Health GmbH, Vienna, Austria; Angai Biotchnology Co., Ltd, Life Health Town, National High-Tech Development Zone, Suzhou, China; Acus Laboratories GmbH, Dueren, Germany; Helmholtz Centre for Infection Research, Braunschweig, Germany; Medical University of Vienna, Department of Laboratory Medicine, Vienna, Austria; Life Science Institute, University of British Columbia, Department of Medical Genetics, Vancouver, British Columbia, Canada; Public Health Agency of Sweden, Solna, Sweden

**Author notes:** Corresponding author and email address Vanessa Monteil and Ali Mirazimi.

**Keywords:** Coronavirus, Virus-host interactions, Screening, Mediator complex, Innate immunity, Interferon

## Abstract

The virus responsible for COVID-19, SARS-CoV-2, continues to spread through the world. The ongoing emergence of new variants with increased viral transmission and immune evasion continue to pose a challenge. Although large-scale genetic screens have identified numerous host factors required for SARS-CoV-2 infection, however the function of these hits remain incompletely understood. In this study, we performed a haploid forward genetic screen in chemically mutagenized mouse embryonic stem cells overexpressing human ACE2 and identified MED13 and DDX60 as important host factors involved in the modulation of SARS-CoV-2 infection. In this study, we have identified and characterized the function of these key element factors for SARS-CoV2 infection. Knockdown of CKM subunits—with the exception of CDK8—or the helicase DDX60 was sufficient to reduce SARS-CoV-2 infection across Vero E6, A549, and Calu-3 cells. During SARS-CoV-2 infection, MED13 was found to function downstream of the JAK/STAT interferon pathway, but also showed another function independently of the interferon response pathway. Surprisingly, while DDX60 is traditionally involved in the interferon response pathway, its knockdown reduces SARS-CoV-2 infection, suggesting DDX60 can promote SARS-CoV-2 infection Interestingly, while inactivation of MED13 or DDX60 markedly reduced SARS-CoV-2 and SARS-CoV infection, it did not affect MERS-CoV. Collectively, these results identify MED13 and DDX60 as critical host determinants for SARS-CoV-2 related-coronaviruses with pandemic potential.

**Impact statement:** Using an unbiased haploid genetic screening approach, this study identifies for the first time MED13 and DDX60 as host factors that influence coronavirus replication through transcriptional and interferon-associated pathways. While DDX60 has previously been linked to antiviral signaling, our findings suggest it can promote coronavirus infection. Most interestingly, the effects of these factors extend beyond SARS-CoV-2 to other coronaviruses.

These findings broaden the current understanding of coronavirus-host interaction by highlighting transcriptional control and interferon-associated pathways as modulators of infection rather than focusing solely on viral entry. The work provides mechanistic insight into how host regulatory networks influence coronavirus replication and suggests potential targets for host-directed therapies with activity against multiple present and future coronavirus threats.

**Data summary:** The authors confirm all supporting data, code and protocols have been provided within the article or through supplementary data files.

**Repositories:** The data generated in this study are provided in the Supplementary Information/Source Data file. Sequencing data are available on NCBI Sequence Read Archive under the accession number BioProject PRJNA1271794.

The next generation sequencing data generated in this study has been deposited in the NCBI Sequence Read Archive (SRA) under accession number SRX29485992, SRX29485993, SRX29485994, SRX29485995, SRX29485996, SRX29485997, SRX29485998, SRX29485999, SRX29486000, SRX29486001, SRX29486002, SRX29486003, SRX29486004, SRX29486005, SRX29486006, SRX29486007, SRX29486010, SRX29486011, SRX29486012, SRX29486013, SRX29486014, SRX29486015, SRX29486016, SRX29486017, SRX29486018, SRX29486019, SRX29486021, SRX29486022, SRX29486023, SRX29486024

## Introduction

The coronavirus disease 2019 (COVID-19) pandemic, caused by severe acute respiratory syndrome coronavirus 2 (SARS-CoV-2), has caused millions of deaths and substantial global economic disruption. SARS-CoV-2 is an enveloped, single-strand, positive-sense RNA virus in the genus *Betacoronavirus* (1, 2). Like other coronaviruses, it exploits host machinery across the infection cycle: entry, genome replication, assembly and release (2, 3). Viral entry essentially depends on Angiotensin-Converting Enzyme 2 (ACE2), acting as the main receptor for SARS-CoV-2, and host proteases such as TMPRSS2 (2, 4). SARS-CoV-2 has continued to evolve during widespread global transmission and Omicron-lineage viruses show broad escape from many neutralizing antibodies and reduced TMPRSS2 usage with reliance on endosomal pathways (5, 6). Worldwide vaccination reduced severe disease but provides incomplete and waning protection against infection, particularly with Omicron-lineage variants (7-9). These features sustain selective pressure on virus-host interactions and motivate systematic mapping of host dependencies to guide host-directed countermeasures (10).

Genome-wide CRISPR knockout screens in human cells identified ACE2 and additional entry and trafficking factors, as well as lipid and glycan pathways and chromatin regulators (3, 11-14). Hit profiles differ by cell type and viral strain (14, 15). Proteomic mapping of SARS-CoV-2 and host protein interactions have further expanded candidate host targets (16). Most of this works has predominantly focused on entry steps, whereas less studies have resolved host factors that modulate viral replication or innate immune responses during infection (17).

Haploid forward genetics enables unbiased identification of recessive host determinants and can capture partial-loss or separation-of-function mutations not readily accessible through CRISPR knockout approaches (18). Our haploid mouse embryonic stem cells (mESCs) provide a powerful platform for unbiased mapping of host factors that control coronavirus infection (19), enabling the investigation of a broad range of functional consequences including loss-of-function, partial loss-of-function or separation-of-function, and gain-of-function mutations. This system has been successfully used for unbiased mammalian genetic screening (3, 20-22).

In this study, we have developed mESCs overexpressing ACE2. By using these cells, we performed an unbiased mutagenesis screen to identify host regulators of SARS-CoV-2 infection and characterized their functional relationship to interferon signaling. We identified MED13, a subunit of the Mediator CDK8 kinase module that regulates RNA polymerase II transcription, and DDX60, an interferon-inducible RNA helicase involved in RIG-I-like receptor (RLR) signaling [20], as cellular determinants that play a role in SARS-CoV-2 infection.

## Methods

### Cells and viruses

Haploid mouse embryonic stem cells (mESCs) AN3-12 are a feeder independent clonal derivative of HMSc2 isolated from mice oocyte and are maintained at Institute of Molecular Biotechnology (IMBA, Austria) (19, 23). AN3-12 cells were validated by Short Tandem Repeat (STR) analysis. Haploid mES cells were maintained in standard ES cell medium (Swedish Veterinary Agency, SVA), supplemented with 10% (v/v) fetal bovine serum (FBS; Gibco), recombinant human Leukemia Inhibitory Factor (LIF; STEMCELL) and β-mercaptoethanol (Sigma-Aldrich). The hACE2-overexpressing haploid mESCs AN3-12 used for haploid screening was obtained from IMBA (Austria). Cells were maintained in ES medium supplemented with 15% (v/v) FBS (Gibco), LIF (STEMCELL) and β-mercaptoethanol (Sigma-Aldrich). A549, Calu-3, and Vero E6 cells were cultured in Dulbecco’s modified Eagle’s medium (DMEM; Gibco), supplemented with 10% (v/v) FBS (Gibco). The SARS-CoV-2 Omicron variant was isolated by the Swedish public health agency (Folkhälsomyndigheten, Stockholm) and propagated in Vero E6 cells. All our experiments involving SARS-CoV-2 were conducted in a biosafety level 3 (BSL-3) laboratory in compliance with the Swedish veterinary agency guidelines (SVA, Uppsala).

### Generation of ACE2 overexpression clone E1

mESC AN3-12 were transduced with two Lentiviral overexpression constructs for human ACE2, pLenti-CMV-ACE2-Myc-DDK-P2A Puro and pLenti-CMV-ACE2-Myc-P2A-Blasti (JLP Health GmbH) and selected with Puromycin and Blasticidin. Single cell derived clones were picked and analyzed for ACE2 expression level. Clone E1 showed strongest ACE2 expression and was used where indicated.

### Chemical mutagenesis of haploid stem cells

Chemical mutagenesis using N-Ethyl-N-nitrosurea (ENU) was performed as described previously (24). Briefly, haploid AN3-12 cells overexpressing human ACE2, were treated, in suspension and under constant agitation, with 0.1 mg/ml ENU in full medium while for 2 h. Cells were then washed 5 times and transferred to a culture dish. Cells were left to recover for 48 h. Cells were then detached and singled using Trypsin/EDTA before being frozen in 10% DMSO, 40% FBS and 50% full medium.

### Chemically mutagenized haploid mouse stem cells screening

Approximately 50 million mutagenized haploid mESCs were seeded in 150 mm cell culture dishes 1 day prior to infection. Attached cells were infected with SARS-CoV-2 at a multiplicity of infection (MOI) of 5 in 8 ml of embryonic stem cell (ES) medium supplemented with 2% fetal bovine serum (FBS). Following a 1-hour incubation at 37 °C in 5% CO_2_, the viral inoculum was retained, and 17 ml of fresh ES medium containing 2% FBS was added to each dish. Culture was then maintained at 37 °C in 5% CO_2_. After 14 days of incubation, cell colonies showing resistance to SARS-CoV-2 induced cytopathic effects (CPE) emerged and were individually isolated and expanded. Putative resistance cells and control cells were seeded at 5 × 10^4^ cells per well in ES medium supplemented with 15% FBS for 24 h, after which the cell were infected with SARS-CoV-2 at an MOI of 5. The CPE was monitored by microscopy for 7 days. All clones that survived infection were subjected to DNA extraction using the Gentra Puregene tissue kit (Qiagen) according to manufacturer’s instructions.

### Whole exome sequencing and screening analysis

Paired end, 150 bp whole exome sequencing was performed on an Illumina Novaseq 6000 instrument after precapture-barcoding and exome capture with the Agilent SureSelect Mouse All Exon kit. For data analysis, raw reads were aligned to the reference genome mm9. Variants were identified and annotated using GATK and snpEff. SARS-CoV2 resistance causing alterations were identified by allelism only considering variants with moderate or high effect on protein and a read coverage >20.

### Statistical Analysis of WES data for SARS-COV-2 resistance screening

The total number of mutations found in all resistant clones isolated and sequenced from the resistance screens was used to determine an average mutation rate. This only considered mutations classified snpEFF high and moderate. The probability to hit each individual gene was calculated from the mutation rate and the genes cds length relative to the total exome length.

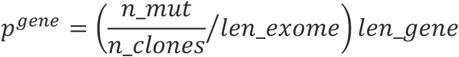

The probability to hit the gene multiple times (e.g. in 4 out of 12 clones) was calculated for each gene using cumulative binomial distribution function.

### *Med13* KO generation in mESCs AN3-12 ACE2 overexpression clone E1

*Med13* targeting sgRNA were selected using VBC-score, www.vbc-score.org (25). sgRNA was cloned into plasmids to express sgRNA, Cas9 and Neomycin resistance as previously described (26). Briefly, oligos CACCGTCACCGGAAACAGAATAG and AAACCTATTCTGTTTCCGGTGAC were annealed, phosphorylated and inserted into Esp3I cloning sites upstream of sgRNA scaffold. 1 × 10^5^ haploid mESCs per well were seeded in a 24-well plate and transfected with 500 ng plasmid 4 h post seeding. 24 h post transfection 500 ug/mL G418 was added to the medium. After 5 days cells were seeded at single cell density and colonies were picked after 12 days. gDNA was extracted and the *Med13* locus genotyped using PCR primers caggccgacttgacaggaat, ACAGCAGAAGCACATTGGGA, Sanger Seq primer cgacttgacaggaattaagtgg.

### *MED13* KO generation in A549 cells

*MED13* targeting sgRNA were selected using VBC-score, www.vbc-score.org (25). sgRNA were cloned into lentiviral Plasmid expressing sgRNA, Cas9 and Puromycin resistance as previously described (26). Briefly, oligos CACCGCTGCAGTAGAAGTTCTTGT and AAACACAAGAACTTCTACTGCAGC were annealed, phosphorylated and inserted into Esp3I cloning sites upstream of sgRNA scaffold. A549 cells were transduced with lentiviral vectors and selected with 1 ug/mL puromycin 24 h post transduction. After 5 days cells were seeded at single cell density and colonies picked after 2 weeks. gDNA was extracted and *MED13* locus genotyped using PCR primers TTTGGACTAAATGGCACTCTCACA, AGGCAGAGCATGCCAAAGTCA, Sanger Seq primer ggacaggcattcaagatgtctg.

### *DDX60* KI generation in mESCs AN3-12 ACE2 overexpression clone E1

*DDX60* Tyr1526>His mutant GAGTAC>GAACAC was generated using co-transfection of HDR donor template and plasmid expressing Cas9 NeoR and sgRNA. sgRNA gcttgcgtattgtactcaagc, single stranded oligo DNA nucleotide repair template CAATCGGGATACCATTAGCAGAAAGGAAGCGAAGTCTTCTGTCACTTGCGTATTGTGTTCAAGCAGGGCGTGACTGAAGTCCTCGGGAAGGTCATCAAGGAACACCTACAA, PCR primer FW AGACCAGTGGCATTTGAGGG, RV primer ATTGTTCTCAGGCAGGCTGT, and Sanger Seq primer caagaagtgaagaaaatgcagc. mESCs E1 cells were seeded at 10% confluency in a 6-well plate. For transfection 100 ul Opti-MEM containing 2 ug Cas9-NeoR-sgRNA plasmid and 300 ng HDR donor template was mixed with 100 ul Opti-MEM (Thermo Fisher) containing 7 ul Lipofectamine (Invitrogen) and added to the cells. 24 h after transfection G418 (Invivogen ant-gn-1) is added for selection at 500 ug/ml. 5 days after Selection single cell derived clones are expanded and genotyped.

### Reverse siRNA transfection

Reverse transfection of wild-type A549 and Vero E6 cells was performed using Lipofectamine RNAiMAX (Thermo Fisher Scientific) and siRNAs targeting *MED13* (On-TARGETplus SMARTpool Human *Med13*; Dharmacon). Briefly, 1 µl siRNA (10 µM) and 3 µl Lipofectamine RNAiMAX were each diluted in 50 µl Opti-MEM Reduced Serum Medium (Gibco), combined by gentle pipetting, and incubated at room temperature for 5 minutes to allow complex formation. The resulting transfection complexes were dispensed into 48-well plates in accordance with the manufacturer’s instructions.

Cells were trypsinized, resuspended in antibiotic-free complete medium, and seeded directly onto the siRNA-lipid complexes at a density of 5 × 10^4^ cells per well. Plates were incubated at 37 °C in 5% CO_2_ for 48 hours prior to infection or analysis.

### Ruxolitinib and IFN-β treatment

A549, Calu-3, and Vero E6 cells were seeded at a density of 5 × 10^4^ cells per well in 48-well plates. At 24 h post seedings, cells were treated with 10 µM of ruxolitinib or 500 IU for 16 h prior to infection. Cell numbers were determined to achieve a MOI of 0.1. The virus inoculum was prepared in DMEM containing 10% FBS and supplemented with the same concentrations of the compounds. Cells were infected for 1 h, washed once with PBS, and fresh DMEM containing 10% FBS and the same concentrations of the compounds was added to each well.

### Infection of A549, Vero E6, and Calu-3 cells

At 48 hours post-transfection, cells were infected with SARS-CoV-2 Omicron variant at a MOI of 0.1 in DMEM and incubated for 24 hours post-infection. Cells were then washed three times with PBS and lysed with TRI Reagent for subsequent analysis by qRT-PCR.

### qRT-PCR

All RNA was extracted using TRI Reagent (Sigma-Aldrich; T9424) and the Direct-zol RNA Extraction Kit (Zymo Research) following the manufacturer’s instructions. Quantitative real-time PCR (qRT-PCR) was performed using the TaqMan Fast Virus 1-Step Master Mix (Thermo Fisher) on a 7500 Real-Time PCR System (Applied Biosystems). SARS-CoV-2 E-gene primers and probe were used as previously described (27). RNaseP RNA served as an endogenous control for normalization. Relative viral RNA levels were calculated using the ΔΔCt method.

### Western blot

Cell pellets were lysed in buffer containing 10 mM Tris, 150 mM NaCl, 0.5% SDS, and 1% Triton X-100 in PBS (pH 7.5; Karolinska Institutet) and mixed with LDS Sample Buffer (NuPAGE; Invitrogen) supplemented with 10% β-mercaptoethanol (Sigma-Aldrich). Lysates were heated at 95 °C for 20 minutes. Protein concentrations were determined using the Pierce 660 nm Protein Assay Reagent (Thermo Fisher Scientific) in combination with the Ionic Detergent Compatibility Reagent (Thermo Fisher Scientific).

Equal amounts of proteins (20 μg per sample) were loaded onto a 4-12% Criterion Bis-Tris gel (Bio-Rad) and transferred to PVDF membranes. Membranes were blocked with 5% (w/v) non-fat dry milk in PBS containing 0.1% Tween-20 (PBS-T) for 1 hour at room temperature, followed by incubation with primary antibodies diluted in 5% milk in PBS-T while rocking for 1 hour.

## Results

### HAPLOID STEM CELL SCREEN IDENTIFIES MED13 AND DDX60 AS POTENTIAL FACTORS INVOLVED IN SARS-COV-2 INFECTION

To uncover cellular factors required for SARS-CoV-2 virus infection, we performed a genome-wide SARS-CoV-2 resistance screen in haploid mouse embryonic stem cells (mESCs). Therefore, mESCs were engineered to express human ACE2 (hACE2) to allow SARS-CoV-2 entry and replication. A hACE2-mESC population was mutagenized with N-ethyl-N-nitrosourea (ENU) to generate a haploid cell library carrying largely random point mutations. Generating haploid screening libraries through chemical mutagenesis enabled interrogation of diverse functional consequences, including complete loss-of-function, hypomorphic, and separation-of-function mutation. Challenged with the SARS-CoV-2 Omicron variant at a cytopathic MOI of 5, colonies resistant to infection were isolated from the cell libraries and expanded. The surviving clones were analyzed by deep sequencing and bioinformatic mapping to identify candidate genes whose mutation conferred resistance (Fig. 1A).

**Figure 1.**
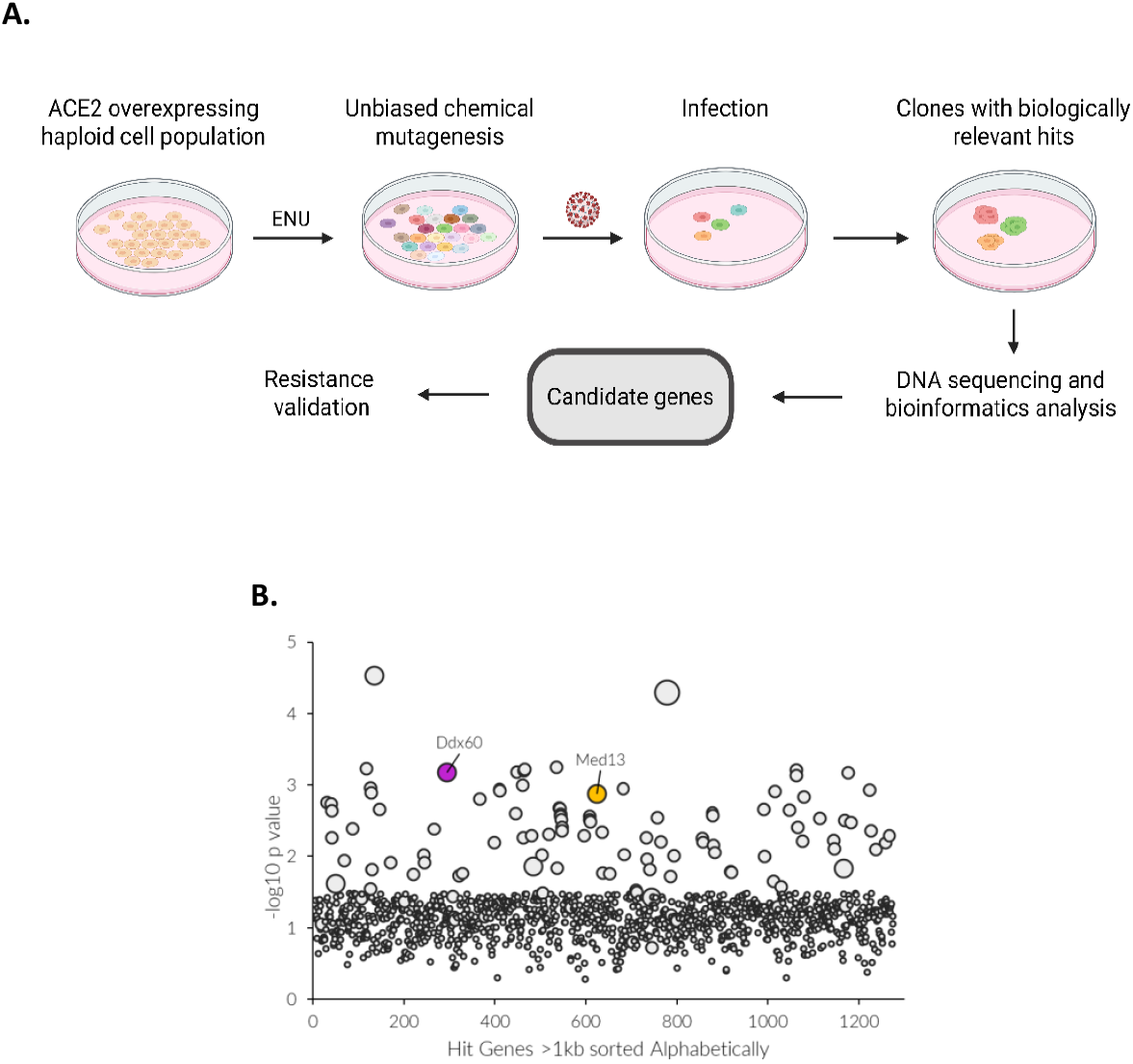
Haploid cells screening identifies *med13* and *ddx60* as host factors involved in SARS-CoV-2 infection. (A) Schematic illustration of the chemical mutagenesis-haploid screening workflow. (B) Chemical mutagenesis haploid screen identifying genes associated with resistance to SARS-CoV-2 infection. Bubble plot of candidate genes identified in the screen. Each bubble represents an individual gene, with bubble size corresponding to the number of unique mutations detected per gene. The y-axis indicates the –log of the P value of total mutations per gene compared with control, and genes are arranged alphabetically along the x-axis

Genome-wide analysis of enriched mutations revealed several candidate hits. Multiple resistant clones carried independent mutations in *Ddx60* and *Med13*, which prompted further validation (Fig. 1B and Supp. Table 1). The enrichment of these loci suggested that they support one or more steps of the SARS-CoV-2 life cycle. Med13, both missense and full-loss alterations were identified, while all three *Ddx60* alleles are missense mutations (Supp. Table 1).

To validate the screen results, we generated independent hACE2-mESC clones carrying either a *Med13* knockout (KO) or a knock-in modification in *Ddx60*. Hereafter, these clones are referred to as KO cells. These KO cells were infected with SARS-CoV-2 at an MOI of 5 and relative viral RNA amount inside the cells was measured by qRT-PCR 24 hours post-infection (24hpi). Viral mRNA in *Med13* KO clones was markedly reduced compared to control cells, whereas the *Ddx60* KO clone showed no significant change (Supp. Fig.1C). To confirm the observed decrease in SARS-CoV-2 infection in *Med13* KO cells is due to the disruption of the *Med13* gene and not because of the loss of hACE2 expression during the generation of the KO cells, hACE2 protein levels in control and KO cells were assessed by western-blot. *Med13* KO cells exhibit a loss of hACE2 protein compared to the control, whereas *Ddx60* KO cells show higher hACE2 expression (Supp. Fig. 1D). Thus, the reduced viral RNA levels observed in *Med13* KO cells (Supp. Fig. 1C) could be attributed to the absence of hACE2 at the cell surface. Likewise, a potentially reduced infection level in *Ddx60* KO might be masked by increased ACE2 levels (Supp. Fig. 1D) and for both KO lines this result cannot be interpreted as an effect of the KO itself.

### MED13 AND DDX60 ARE KEY HOST FACTORS FOR SARS-COV-2 IN DIPLOID CELLS

To validate the hits highlighted by the haploid screen while avoiding the bias of the ACE2 expression, we decided to use cell models naturally expressing ACE2. Thus, we performed siRNA-mediated knockdown in diploid cell lines, Vero E6 and Calu-3, and quantified viral mRNA levels at 24hpi. In Vero E6 (Fig. 2A), *MED13* knockdown resulted in a strong reduction of viral RNA (around 55% decrease; ***P < 0.001), whereas *DDX60* knockdown led to a moderate but significant reduction in viral RNA levels (> 25% decrease; *P < 0.05). In Calu-3 (Fig. 2B), the knockdown of *MED13* almost abolished viral RNA (> 90% decrease; ***P < 0.0001) and *DDX60* knockdown similarly led to near-complete loss of viral RNA (***P < 0.0001). Together, these results confirm MED13 and DDX60 as host factors important for SARS-CoV-2 infection while their infection promoting effects seem to be cell-type-dependent.

**Figure 2.**
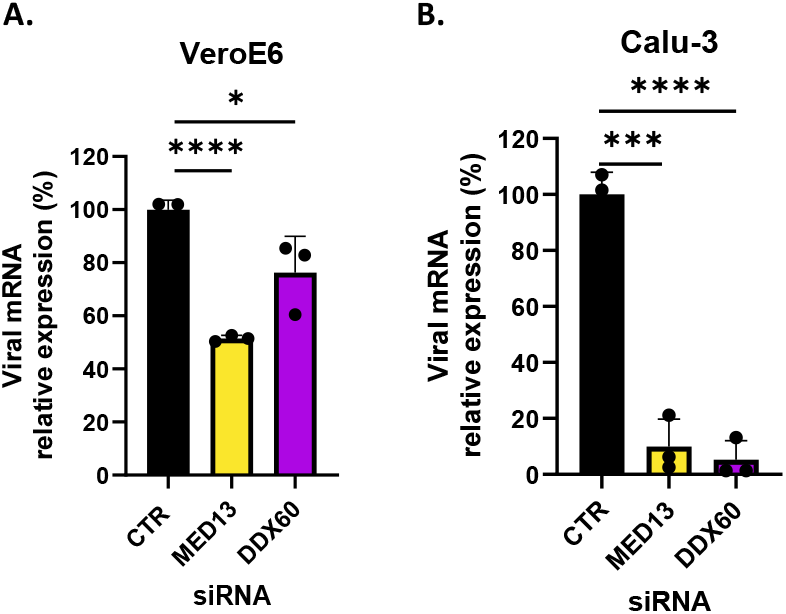
Validation of *MED13* and *DDX60* as host genes involved in SARS-CoV-2 infection in diploid cells. The knockdown of *MED13* and *DDX60* impairs SARS-CoV-2 infection in diploid cells. Control, *MED13*-knockdown and *DDX60*-knockdown (A) Vero E6 and (B) Calu-3 cells were infected with SARS-CoV-2 (MOI = 0.1) and harvested 24 hpi. Data are shown as percentage of viral mRNA expression relative to control cells (n = 3 independent experiments, mean ± SD, Student’s *t*-test (\**p* < 0.05, \*\*\**p* < 0.001, \*\*\*\**p* < 0.0001).

### MED13 ACTS THROUGH THE MEDIATOR CDK8 KINASE MODULE, BUT CDK8 IS DISPENSABLE

The mediator complex is a large multiprotein complex of almost 30 proteins that connect transcription factors to RNA polymerase II and regulate transcription. A part of the Mediator complex is the CDK8 kinase module (CKM). The CKM regulates RNA polymerase II transcription (28). This module is a complex of several proteins: MED12, MED13, CDK8, and Cyclin C (CCNC) (Fig. 3A). To assess the role of MED13 on SARS-CoV-2 as a member of the CKM, we asked whether other subunits of the module could also affect SARS-CoV-2 infection. In Vero E6 cells, the knockdown of *MED13, MED12*, or *CCNC* significantly reduced viral RNA, whereas the knockdown of *CDK8* had little or no effect (Fig. 3B). In A549 cells, the knockdown of *MED13* and *MED12* decreased viral RNA, while *CDK8* and *CCNC* did not significantly affect infection (Fig. 3B).

**Figure 3.**
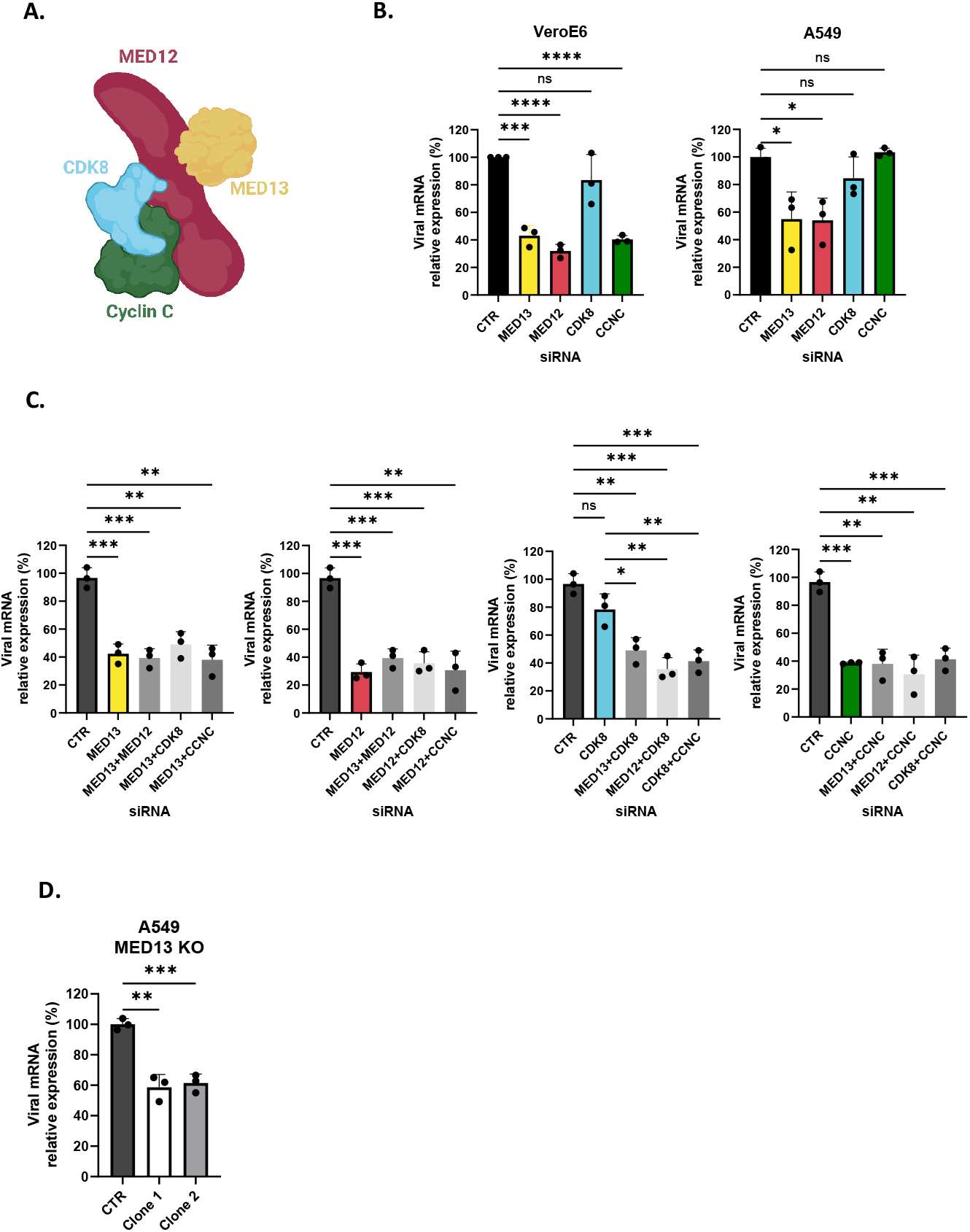
MED13 is involved in SARS-CoV-2 infection through the CDK8 kinase module. As components of the CDK8 kinase module (CKM), MED13 and associated subunits were evaluated in SARS-CoV-2 infection. (A) Schematic illustration of the CKM and its subunits. (B) Individual CKM subunits were knocked down in Vero E6 and A549 cells, followed by infection with SARS-CoV-2 (MOI = 0.1). Cells were harvested at 24 hpi. (C) Effects of combined knockdown of CKM subunits on SARS-CoV-2 in Vero E6 cells. Cells were infected at MOI = 0.1 and harvested at 24 hpi. (D) Validation of *MED13* knockout in A549 cells by quantification of *MED13* mRNA level. Data are shown as percentage of viral mRNA expression relative to control cells (n = 3 independent experiments, mean ± SD, Student’s *t*-test, \**p* < 0.05, \*\**p* < 0.01, \*\*\**p* < 0.001, \*\*\**p* < 0.0001).

We next performed double-knockdowns of CKM subunits in Vero E6 cells to assess the potential cumulative or synergetic effects on SARS-CoV-2 infection (Fig. 3C). Double knockdowns involving *MED13* or *MED12* did not further reduce viral levels compared with the corresponding single knockdowns indicating that these factors likely function within the same pathway to restrict SARS-CoV-2 infection. In contrast, combinations that included *CDK8* (i.e. *MED13*+*CDK8, MED12*+*CDK8, CDK8*+*CCNC*) reduced viral RNA levels relative to *CDK8* knockdown alone, however knockdown of *CDK8* did not enhance the antiviral effect observed upon knockdown of the other CKM subunits individually. These data suggest that CDK8 itself does not play a critical role in SARS-CoV-2 infection and may be dispensable for the CKM associated function, relevant to viral replication in Vero E6 cells. To further confirm the importance of MED13 in SARS-CoV-2 infection, *MED13* knockout A549 cells were generated (Supp. Fig 2) and infected with SARS-CoV-2. Consistent with the knockdown experiments *MED13* knockout resulted in comparable reduction in viral RNA levels (Fig. 3D).

Together, these data highlighted MED13 as an important factor involved in SARS-CoV-2 infection, via its role as a part of the CKM.

### MED13 RESTRICTS SARS-COV-2 INFECTION THROUGH A JAK/STAT DEPENDENT INTERFERON SIGNALING

As shown in Fig. 2B, the knockdown of *DDX60* led to a significant reduction in SARS-CoV-2 infection in Calu-3 cells. DDX60 is an interferon-inducible helicase that participates in RIG-1/MDA5 driven antiviral signaling (29). Based on these findings, we next asked whether the role of MED13 in SARS-CoV-2 infection could intersect with interferon pathways. To this aim, we used Calu-3 cells as they are competent for both IFN production and response (17). To assess the role of interferon signaling, JAK/STAT activation was inhibited using JAK1/2 inhibitor ruxolitinib in control and *MED13* knocked down Calu-3 cells and SARS-CoV-2 infection levels were assessed. As we previously shown (Fig.2B), the knockdown of *MED13* reduced viral infection, but ruxolitinib treatment restored viral mRNA to control levels (Fig. 4A). This recovery of infection indicates that the antiviral effect of *MED13* knockdown is, at least in part, JAK/STAT dependent on Calu-3 cells. In *MED13* knockdown Vero E6 cells and *MED13* knockout A549 clones, according to previous data (Fig. 2A and 3D), the level of infection is significantly reduced, while ruxolitinib treatment does not affect SARS-CoV-2 infection in these cells (Fig. 4B and 4C) contrary to Calu-3 (Fig. 4A). in agreement with the findings of others (4, 30). The lack of rescue is compatible with the weak to lack of interferon production or response typically observed in A549 and Vero E6 cells during SARS-CoV-2 infection (30, 31), implying that while MED13 showed to be involved in the IFN response in Calu-3 cells, another mechanism involving MED13 also affect SARS-CoV-2 infection in an IFN-independent manner.

**Figure 4.**
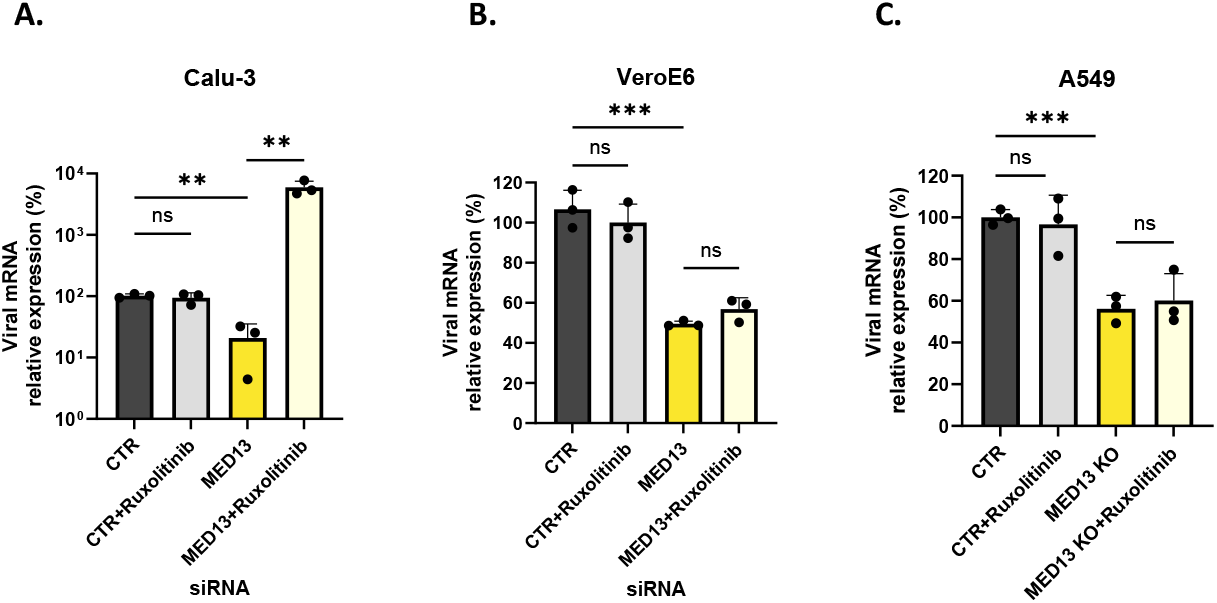
Effect of Ruxolitinib on *MED13* knockdown or knockout cells. The effect of Ruxolitinib on SARS-CoV-2 infection were evaluated in control and *MED13*-deficient cells. Control and *MED13*-knockdown or knockout cells were pretreated with Ruxolitinib (10uM) or DMSO for 24 h, followed by infection with SARS-CoV-2 (MOI = 0.1). (A) Calu-3, (B) Vero E6 cells, (C) A549 cells were harvested 24hpi. Data are presented as the percentage of viral mRNA expression relative to control cells. (n = 3 independent experiments, mean ± SD). Statistical significance was determined using Student’s *t*-test (\**p* < 0.05, \*\**p* < 0.01, \*\*\**p* < 0.001).

### MED13 AND DDX60 EFFECTS IN CALU-3 CELLS DEPEND ON AN INTACT JAK/STAT SIGNALING

To further dissect the contribution of interferon signaling to the antiviral effects mediated by MED13 and DDX60, we examined the impact of JAK/STAT inhibition on SARS-CoV-2 infection in interferon-competent Calu-3 cells. Control cells, as well as cells with *MED13*, or *DDX60* knockdown were pretreated with the JAK1/2 inhibitor ruxolitinib prior to infection, and viral RNA levels were quantified at 24 h post-infection (Fig. 5A). In Calu-3 cells, ruxolitinib treatment alone did not significantly alter viral RNA levels, consistent with previous observations (Fig. 4A; Fig. 5A,). As shown earlier (Fig. 2B), knockdown of MED13 or DDX60 resulted in a marked reduction in viral replication. Notably, ruxolitinib treatment restored viral RNA levels in both MED13-and DDX60-knockdown cells, indicating that MED13 and DDX60 affect SARS-CoV-2 infection via the JAK/STAT signaling pathway in Calu-3 cells (Fig. 5A). Consistent with this, while the knockdown of *MED13* induces the expression of the interferon-stimulated gene *MX1* and *DDX60*, ruxolitinib treatment significantly suppressed the induction of *MX1* and *DDX60* in control as well as *MED13*- and *DDX60*-knockdown cells (Fig. 5B and 5C). Together, these results indicate that MED13 and DDX60 contribute to the interferon-mediated antiviral response to SARS-CoV-2 infection in Calu-3 cells.

**Figure 5.**
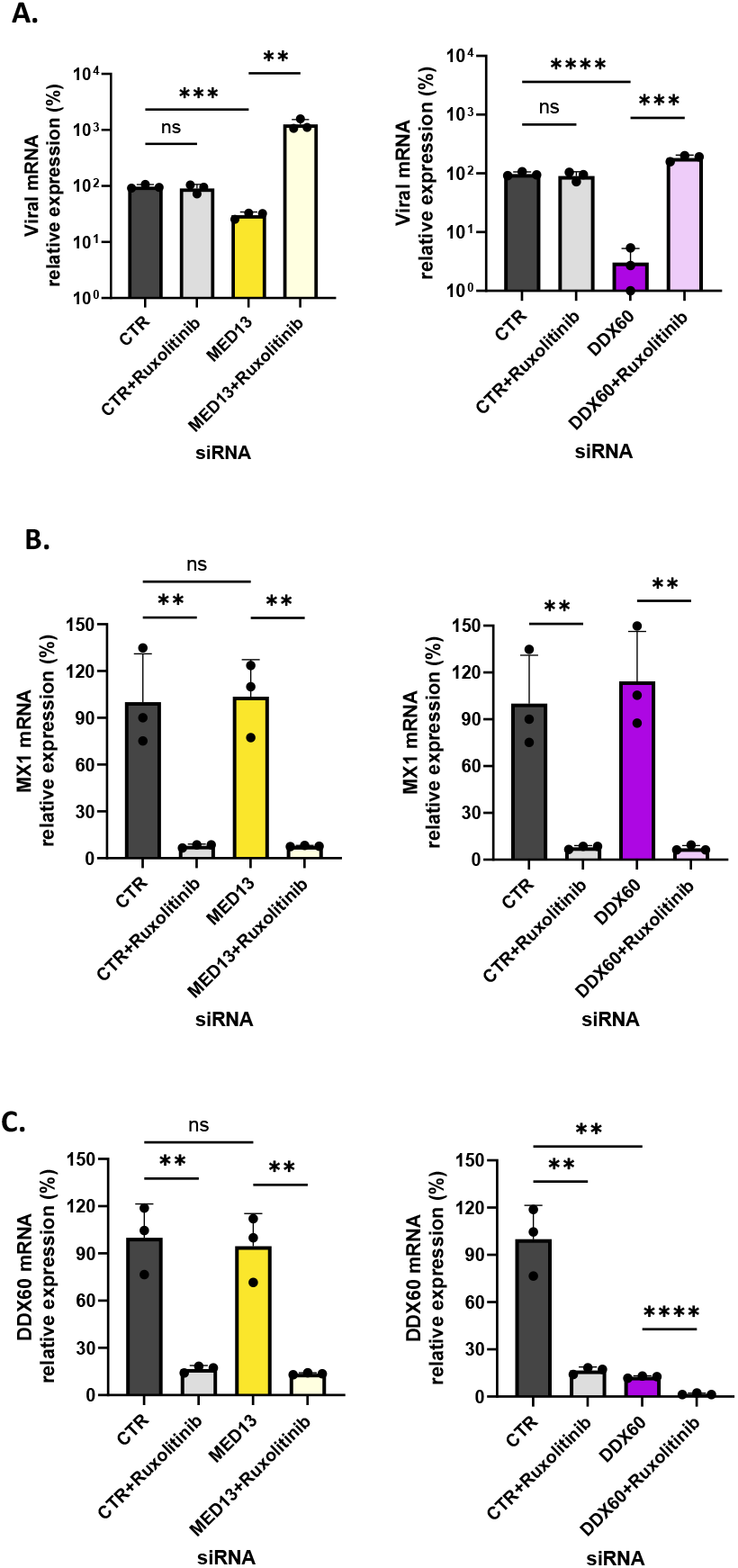
Effect of Ruxolitinib on SARS-CoV-2 replication in *MED13 knockdown* Calu-3 cells. The effect of Ruxolitinib on SARS-CoV-2 Omicron were evaluated in control and *MED13*-knockdown Calu-3 cells. Control and *MED13*-knockdown cells were pretreated with Ruxolitinib (10uM) or DMSO for 24 h, followed by infection with SARS-CoV-2 (MOI = 0.1). Cells were harvested at 24 hpi for analysis. (A) Viral mRNA levels were quantified by real-time RT-PCR and presented as the percentage of viral mRNA expression relative to control cells. (B) *MX1* mRNA levels were quantified by real-time RT-PCR and are presented as the percentage of *MX1* mRNA expression relative to control cells. (C) *DDX60* mRNA levels were quantified by real-time RT-PCR and are presented as the percentage of *DDX60* mRNA expression relative to control cells. Data represent mean ± SD from three independent experiments (n = 3). Statistical significance was determined using Student’s *t*-test (\*\**p*< 0.01, \*\*\**p* < 0.001, \*\*\*\**p* < 0.0001).

### IFN-B SUPPRESSES SARS-COV-2 REPLICATION IN INTERFERON-DEFICIENT VERO E6 CELLS

While ruxolitinib does not restore SARS-CoV-2 infection in *MED13* KD Vero E6 cells (Fig. 4B) contrary to Calu-3 cells (Fig.4A) for which we showed the involvement of the interferon-response pathway, we hypothesized that the differences observed between Calu-3 and Vero E6 were due to the lack of interferon production by the Vero, instead of a lack of interferon response. We tested the interferon-response pathway using exogenous interferon stimulation. Control, *MED13*-, or *DDX60*-KD Vero E6 cells were pretreated with IFN-β prior to infection with SARS-CoV-2 (Fig. 6)

**Figure 6.**
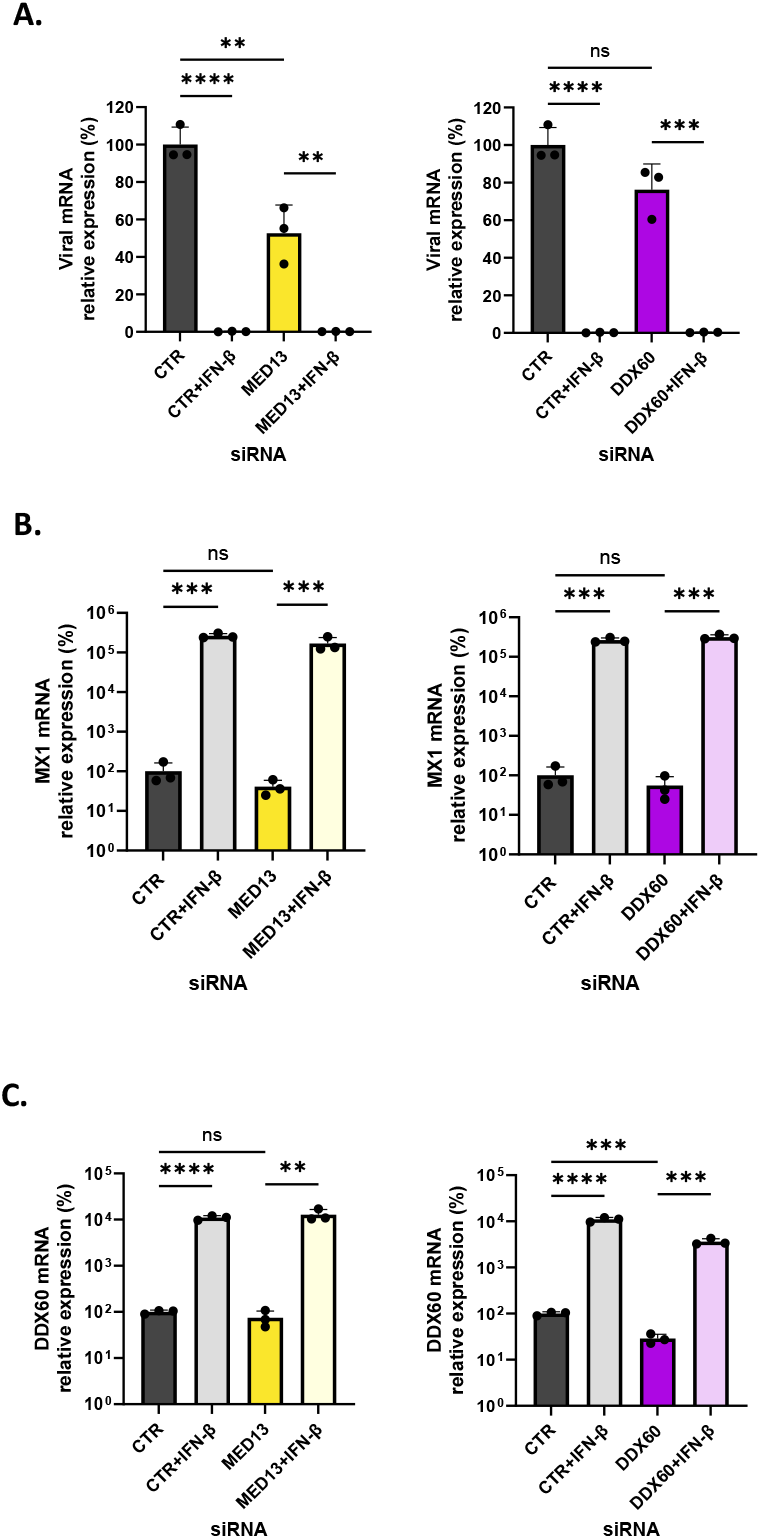
IFN-response pathway is intact in Vero E6 cells. The effect of IFN-β on SARS-CoV-2 Omicron infection were assessed in interferon-deficient Vero E6 cells. (A) Control, *MED13*-knockdown, and *DDX60*-knockdown cells were pretreated with IFN-β (500IU) or DMSO for 24 h, followed by infection with SARS-CoV-2 (MOI = 0.1). Cells were harvested at 24 hpi for analysis. Viral mRNA levels were quantified by real-time RT-PCR and are presented as the percentage of viral mRNA expression relative to control cells. (B) *MX1* mRNA levels were quantified by real-time RT-PCR and are presented as the percentage of *MX1* mRNA expression relative to control cells. (C) *DDX60* mRNA levels were quantified by real-time RT-PCR and are presented as the percentage of *DDX60* mRNA expression relative to control cells. Data represent mean ± SD from three independent experiments (n = 3). Statistical significance was determined using Student’s *t*-test (\*\**p* < 0.01, \*\*\**p* < 0.001, \*\*\*\**p* < 0.0001).

IFN-β pretreatment resulted in a significant reduction of viral mRNA in control cells, confirming the functionality of the interferon signaling pathway in this interferon production-deficient cell line (Fig. 6A and 6B) (30). In *MED13*-, or *DDX60*-KD cells, IFN-β treatment also dramatically decreased viral RNA levels.

As expected, IFN-β pretreatment robustly induced expression of *MX1* across all conditions (Fig. 6B). In addition, *DDX60* mRNA levels were strongly up-regulated following IFN-β stimulation, confirming that downstream of interferon receptor signaling remains intact in Vero E6 cells (Fig. 6C). Together, these results demonstrate that the IFN response signaling in Vero E6 is intact and that the reduction in SARS-CoV-2 infection in *MED13* but also in *DDX60* KD cells are independent of the IFN response pathways, contrary to Calu-3.

### IFN-B ENHANCES SARS-COV-2 REPLICATION IN INTERFERON-RESPONSE DEFICIENT A549 CELLS

We examined the effect of IFN-β treatment in A549 cells, which are known to have impaired interferon response during SARS-CoV-2 infection (4, 32). Wild-type and *MED13*-knockout A549 cells were pretreated with IFN-β prior to infection with SARS-CoV-2 and viral RNA levels were quantified at 24hpi (Fig. 7).

**Figure 7.**
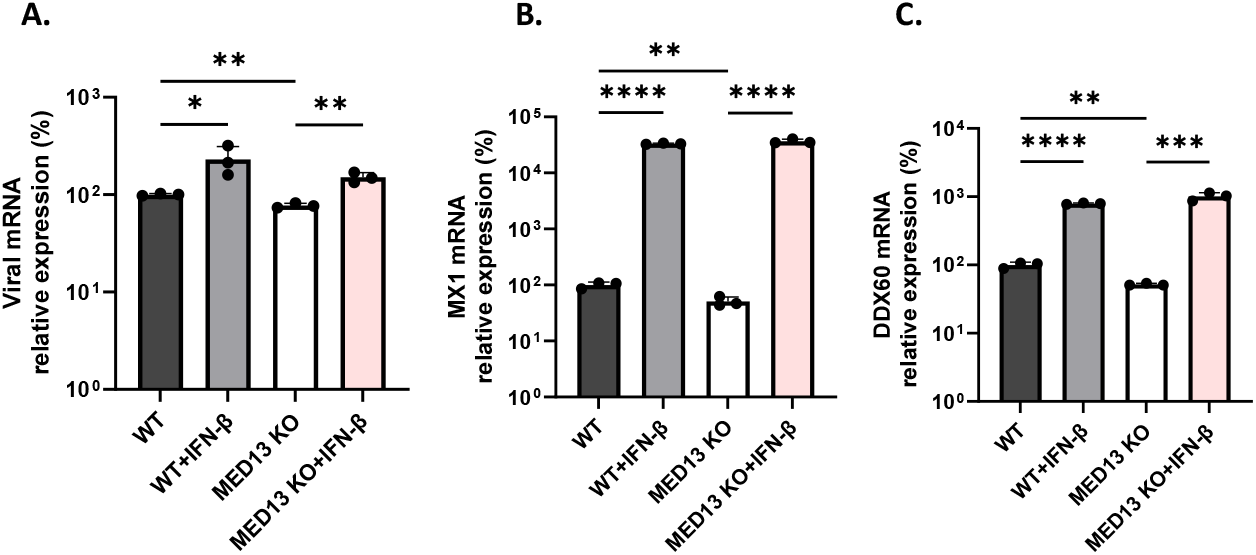
IFN-β induces interferon-stimulated gene expression in A549 cells but fails to restrict SARS-CoV-2 replication. The effect of IFN-β on SARS-CoV-2 infection were assessed in interferon-deficient A549 cells. Wild-type and *MED13* knockout clone cells were pretreated with IFN-β (500IU) or DMSO for 24 h, followed by infection with SARS-CoV-2 (MOI = 0.1). Cells were harvested at 24 h post-infection for analysis. (A) Viral mRNA levels were quantified by real-time RT-PCR and presented as the percentage of viral mRNA expression relative to control cells. (B) *MX1* mRNA levels were quantified by real-time RT-PCR and are presented as the percentage of *MX1* mRNA expression relative to control cells. (C) *DDX60* mRNA levels were quantified by real-time RT-PCR and are presented as the percentage of *DDX60* mRNA expression relative to control cells. Data represent mean ± SD from three independent experiments (n = 3). Statistical significance was determined using Student’s *t*-test (\**p* < 0.05, \*\**p* < 0.01, \*\*\**p* < 0.001, \*\*\*\**p* < 0.0001).

In contrast to Calu-3 and Vero E6 cells, IFN-β pretreatment did not suppress SARS-CoV-2 replication in A549 cells. Surprisingly, IFN-β treatment resulted in a significant increase in viral RNA levels in wild-type A549 cells compared to untreated control (Fig. 7A). A similar increase was observed in *MED13*-knockout cells (Fig 7A), indicating that the promoting effect of IFN-β on viral replication in this cell type occurs independently of *MED13* expression.

Despite the lack of antiviral activity, IFN-β pretreatment robustly induced the expression of the IFN-stimulated genes *MX1* and *DDX60*, in both wild-type and *MED13* knockout A549 cells (Fig. 7B-C). These results demonstrate that A549 cells retain the capacity to respond to exogenous IFN-β, however IFN-β induced signaling is insufficient to restrict SARS-CoV-2 replication in this cellular context.

### MED13 AND DDX60 ARE IMPORTANT FOR SARS-COV INFECTION BUT NOT FOR MERS-COV

As other *betacoronaviruses* are a threat for humans, we decided to assess the possible involvement of these proteins in the SARS-CoV-2 closely related SARS-CoV and in the more phylogenetically distant MERS-CoV infection. Calu-3 cells knocked down for *MED13* and *DDX60* were infected with either SARS-CoV or MERS-CoV at a MOI of 0.01. 24hpi, the level of viral RNA into cells were measured by RT-qPCR. As shown in Figure 8, the knock down of *MED13* as well as *DDX60* affect SARS-CoV infection (Fig.8A), while not affecting MERS-CoV infection (Fig.8B). These data highlight the importance of MED13 and DDX60 for SARS-CoV-2 related viruses while other emerging coronaviruses are not affected by these factors.

**Figure 8.**
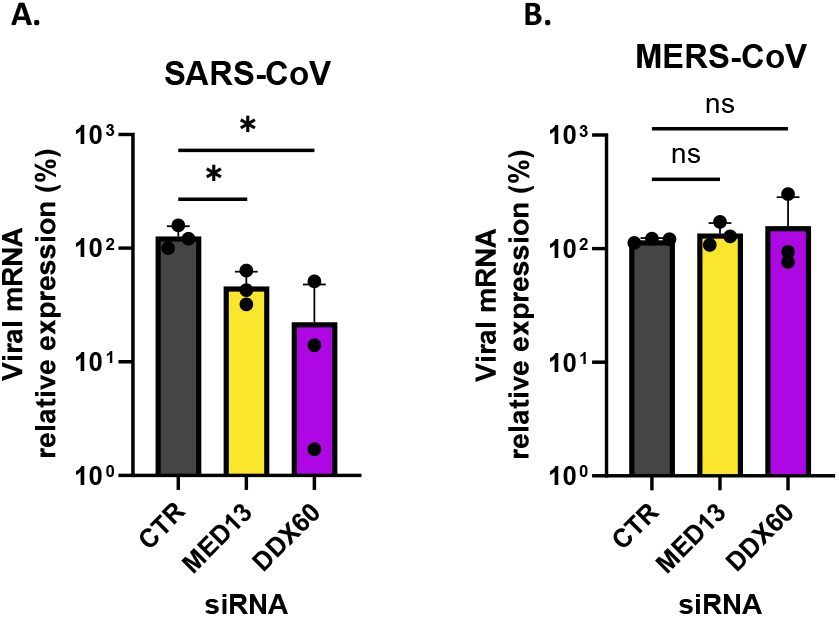
Knockdown of *MED13* and *DDX60* reduces SARS-CoV but not MERS-CoV infection in Calu-3 cells. Control siRNA, *MED13*-siRNA, and *DDX60*-siRNA tranfected Calu-3 cells were infected with SARS-CoV or MERS-CoV (MOI = 0.01) and harvested at 24 hpi. Viral RNA levels were quantified by real-time RT-PCR. Data represent mean ± SD from three independent experiments (n = 3). Statistical significance was determined using Student’s *t*-test (\**p* < 0.05).

## Discussion

Using the hACE2-expressing haploid cells screening, we identified MED13 and DDX60 as candidate host factors influencing in SARS-CoV-2 infection (Fig. 1B). Although validation of these hits in the engineered haploid cells was complicated by clonal variation in ACE2 expression, the principal SARS-CoV-2 entry receptor, ACE2 (Supp. Fig. 1D), the use of diploid cell lines consistently confirmed both MED13 and DDX60 as important factors for SARS-CoV-2 infection (Fig. 2A and 2B).

Our data suggests that the effect of MED13 in SARS-CoV-2 infection relates to its function within the mediator CDK8 kinase module (CKM). Single- and double-knockdown experiments demonstrated that knockdown of the CKM scaffold subunits related genes *MED13, MED12*, or *CCNC* significantly reduced viral infection, whereas knockdown of *CDK8* alone had no measurable effect (Fig. 3B). These findings suggest that CDK8 catalytic activity may be dispensable for SARS-CoV-2 infection under our experiment conditions. This observation is consistent with the established architecture of the CKM (Fig 3A), in which MED13 and MED12 function as adaptors proteins that bridge the CCNC-CDK8 to the core Mediator complex to regulate stimulus-responsive transcription (33). The absence of additive effects in double-knockdowns (Fig. 3C) further supports the conclusion that these subunits act within a shared, module-dependent pathway rather than as independent factors.

Given the central role of the Mediator complex as a transcriptional coactivator bridging transcription factors to RNA polymerase II (28, 33), and regulating a large fraction of cellular gene expression, any effect of MED13 on SARS-CoV-2 replication is likely indirect and mediated through altered host transcriptional programs. Because SARS-CoV-2 is a positive-sense RNA virus that does not generate a DNA intermediate, Mediator-dependent effects on viral replication most likely arise from alterations in host transcriptional pathways, such as innate immune signaling or metabolic pathways relevant to viral replication.

Distinct mechanisms of action were observed across different cell types. In interferon-competent Calu-3 cells, knockdown of either *MED13* or *DDX60* markedly reduced SARS-CoV-2 infection. In contrast, Vero E6 cells, which lack type I interferon production due to a large genomic deletion (34), and A549 cells, which display impaired interferon responses due to oncogenic KRAS signaling (35), also showed significant reductions in viral replication following *MED13* knockdown or knockout despite minimal interferon activity. These observations indicate that MED13 supports SARS-CoV-2 infection through an interferon-dependent and interferon-independent mechanism, depending on cell types.

In Calu-3 cells where SARS-CoV-2 elicits an MDA5-dependent type I and III interferon response (17), knockdown of *MED13* or *DDX60* markedly reduced viral infection (Fig. 4A). Ruxolitinib, a clinically approved JAK1/2 inhibitor that has been investigated as a therapeutic approach for COVID-19 due to its ability to dampen excessive cytokine signaling (16, 36, 37), restored viral RNA levels in *MED13*- or *DDX60*-knockdown Calu-3 cells, suggesting that their effects are at least partly depend on intact JAK/STAT-mediated interferon signaling. By contrast, ruxolitinib did not modify the reduced infection observed in *MED13*-knockdown Vero E6 or knockout A549 cell clones (Fig. 4B and 4C), consistent with the weak or absent interferon competence of these cell lines during SARS-CoV-2 infection (4).

Our data further distinguishes the effects of endogenous interferon signaling triggered by viral infection from response to exogenous interferon stimulation. Although Vero E6 cells are not capable to endogenously produce type I IFNs, treatment with exogenous type I IFN robustly induced IFN-stimulated genes, including *MX1* and *DDX60*, and suppressed SARS-CoV-2 infection. These findings confirm that downstream interferon pathways remain intact in Vero E6 cells despite defective interferon production.

In contrast, exogenous IFN-β treatment failed to restrict SARS-CoV-2 replication in A549 cells despite strong induction of interferon-stimulated genes. IFN-β treatment even modestly enhanced viral RNA levels. In line with this, A549 cells are known to carry an oncogenic KRAS mutation that has been reported to modulate downstream interferon signaling, failing to show a reduction in viral replication (35).

Taken together, the differences observed between the different cell lines in our study highlight the importance of using several models to assess the role of proteins in viral infection, with physiologically relevant systems such as organoid models representing an important future direction.

Mechanistically, DDX60 is an interferon-inducible RNA helicase previously reported to promote RIG-1- and MDA5-dependent antiviral signaling and viral RNA degradation (29, 38). Although DDX60 is an interferon-stimulated gene implicated in antiviral response, it was unexpected that its knockdown increases basal Interferon Stimulated Gene expression (MX1) and renders cells more resistant to SARS-CoV-2 infection (Fig 2), an effect that could be reversed by Ruxolitinib (Fig. 5A). A possible explanation is that SARS-CoV-2 uses DDX60 for its own sake. Thus, DDX60 has been reported to be hijacked by Crimean-Congo hemorrhagic fever virus to promote viral replication through G-quadruplex (G4) unwinding, facilitating viral RNA metabolism (39). As SARS-CoV-2 genomic RNA has been shown to contain G-quadruplex structures (40-42), it is one speculative possibility that DDX60 may similarly enhance viral replication by resolving these secondary RNA structures. This hypothesis remains speculative but may help reconcile the observed reduction in viral replication following DDX60-knockdown and requires further investigation.

Notably, MED13 and DDX60 are also important for the closely related SARS-CoV but not for the other *betacoronaviruses* MERS-CoV that is more distant on the phylogenetic tree.

In the future, it may be important to uncover the other mechanisms by which MED13 affects SARS-CoV-2/SARS-CoV infection, using more relevant models like organoids. Our study is the second one highlighting DDX60 as a cellular factor diverted from its antiviral role to promote infection, yet further mechanistic studies will be required to explain the mechanisms involved.

In summary, our results identify two host proteins influencing SARS-CoV-2 infection: a CKM-dependent transcription pathway centered on MED13, and more importantly, DDX60, while traditionally involved in the IFN-related antiviral pathway, can function as a negative regulator of the IFN system. These data open a potential therapeutic avenue: targeting negative regulators of interferon signaling may induce tonic interferon-stimulated gene expression and thereby enhance baseline antiviral protection. Such an approach might be more targeted and potentially more sensitive than recombinant interferon therapy, as it leverages the natural expression of type I and type III interferons.

## Author statements

### Author contributions

This study was conceived and designed by H.K., V.M., A.M., M.H.. H.K., V.M. wrote the manuscript. H.K. and T.W. performed all the experiments involving viruses/western blot/qRT-PCR/analysis. M.H., M.D., G.M., A.M., A.B., M.K. developed the chemical mutagenesis haploid stem cells library screening used in this study and prepared the KO A549 cells. M.D., G.M. and M.H. developed and analyzed the data from the mutagenesis haploid screening. A.M., J.M.P., F.W., L.J., V.M., M.H. edited the manuscript.

### Ethical approval

Not applicable.

### Consent for publication

Not applicable.

## Acknowledgements

We acknowledge Dr Max J. Kellner for providing feedback on the manuscript.

