## Supplementary figures/table for "Identification of MED13 and DDX60 as critical host factors for SARS-CoV-2 infections"

Supplementary Figure 1

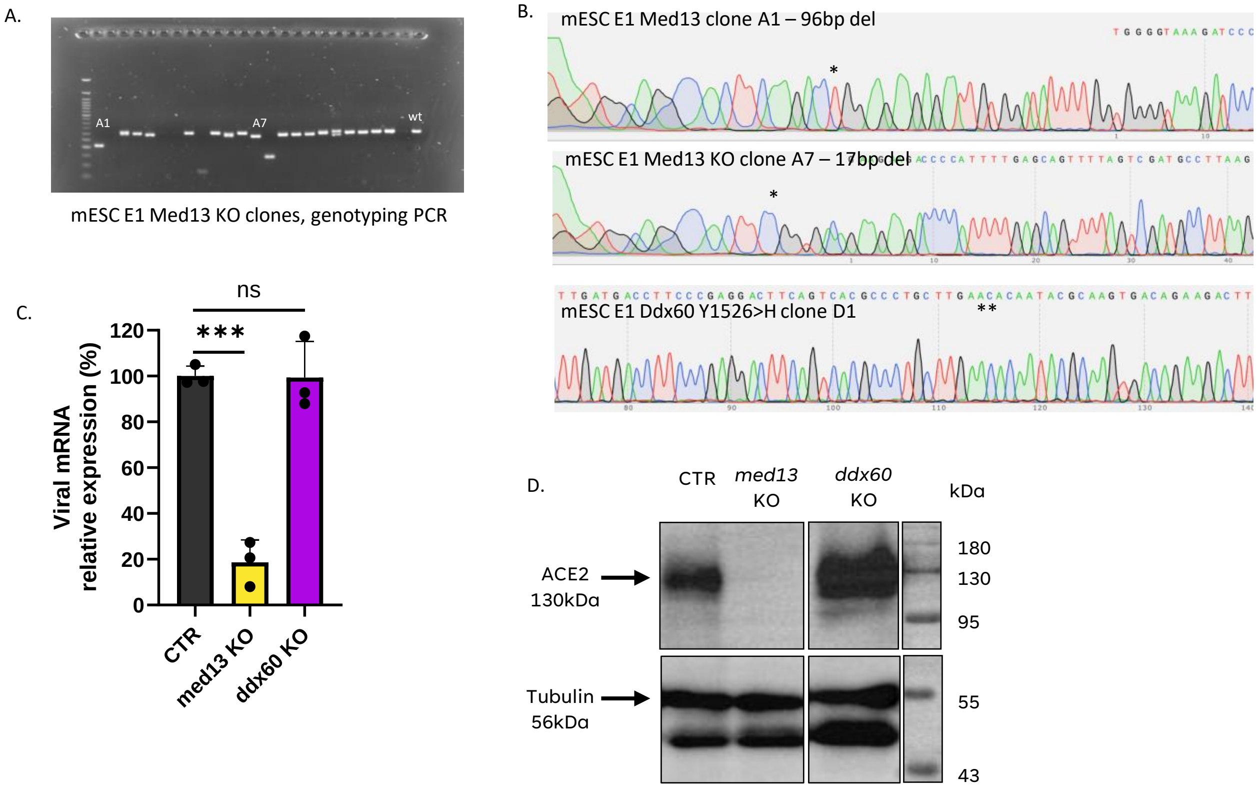

Supplementary Fig 1.

A) Agarose gel of genotyping PCR products on edited Med13 locus in mESC clones. B) Sanger Sequencing tracks of selected clones, positions of genomic DNA edits are marked with an asterisk. (C) Control (CTR), *med13*-knockout, and *ddx60*-knockout haploid cells were infected with SARS-CoV-2 (MOI = 5). Cells were harvested 24hpi, and relative SARS-CoV-2 mRNA levels were quantified. Data are shown as percentage of viral mRNA expression relative to control cells (n = 3 independent experiments, mean ± SD, Student's *t*-test, ***$p$ < 0.001). (D) Western blot analysis of hACE2 protein levels in control and knockout haploid cell clones.

Supplementary Figure 2

A.

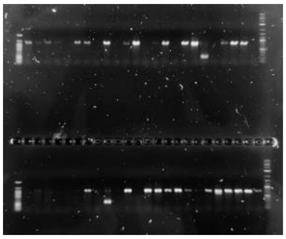

A549 MED13 KO clones, genotyping PCR

B.

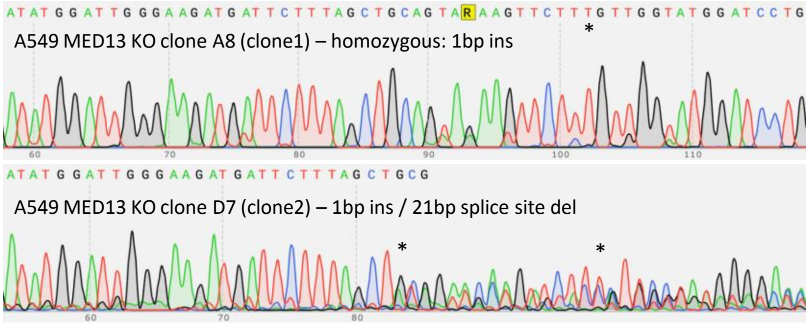

Supplementary Figure 2.

Agarose gel of genotyping PCR products on edited Med13 locus in A549 clones. b) Sanger Sequencing tracks of selected clones, positions of genomic DNA edits are marked with an asterisk.

Supplementary Table 1: List of identified genetic alterations in the selected candidate genes.

| sampleID | chrom | pos | REF base | ALT base | Gene | Mutation consequence |
|---|---|---|---|---|---|---|
| CoV2-Om_N5 | chr11 | 86114281 | A | T | Med13 | Leu919* |
| CoV2-Om_N13 | chr11 | 86111780 | A | C | Med13 | splice donor variant |
| CoV2-Om_N22 | chr11 | 86100392 | C | T | Med13 | Gly1460Asp |
| CoV2-UK_14.1 | chr11 | 86170984 | C | T | Med13 | Cys15Tyr |
| CoV2_Om_N11 | chr8 | 64501995 | T | C | Ddx60 | Tyr1526His |
| CoV2-UK_14 | chr8 | 64500108 | G | A | Ddx60 | Val1477Ile |
| CoV2-UK_7 | chr8 | 64501959 | G | T | Ddx60 | Asp1514Tyr |
